# Measurement of Kinetics of Hammerhead Ribozyme Cleavage Reactions using Toehold Mediated Strand Displacement

**DOI:** 10.1101/2020.09.19.304931

**Authors:** Jay Bhakti Kapadia, Nawwaf Kharma, Alen Nellikulam Davis, Nicolas Kamel, Jonathan Perreault

## Abstract

This paper presents a probe comprising a fluorophore and a quencher, enabling measurement of hammerhead ribozyme cleavage reactions, without labeled RNA molecules, regular sampling or use of polyacrylamide gels. The probe is made of two DNA strands; one strand is labelled with a fluorophore at its 5’-end, while the other strand is labelled with a quencher at its 3’-end. These two DNA strands are perfectly complementary, but with a 3’-overhang of the fluorophore strand. These unpaired nucleotides act as a toehold, which is utilized by a detached cleaved fragment (coming from a self-cleaving hammerhead ribozyme) as the starting point for a strand displacement reaction. This reaction causes the separation of the fluorophore strand from the quencher strand, culminating in fluorescence, detectable in a plate reader. Notably, the emitted fluorescence is *proportional* to the amount of detached cleaved-off RNAs, displacing the DNA quencher strand. This method can replace or complement radio-hazardous unstable ^32^P as a method of measurement of the kinetics of ribozyme cleavage reactions; it also eliminates the need for polyacrylamide gels, for the same purpose. Critically, this method allows to distinguish between the total amount of cleaved ribozymes and the amount of *detached* fragments, resulting from that cleavage reaction.

## INTRODUCTION

Nucleic acid strand displacement reactions make DNA and RNA into powerful tools for the design of various molecular motors (Yurke et al. 2000), biosensors and amplifiers, as well as a means of molecular computation akin to digital circuits and neural network (Qian and Winfree 2011; Qian et al. 2011; Wang et al. 2018). DNA is often the material of choice for building nano circuits, amplifiers and molecular probes, often preferred over RNA due to DNA’s robustness and ease of production and manipulation (Zhang and Winfree 2009). Toehold mediated strand displacement reactions (TMSDRs) are widely used to determine single nucleotide polymorphisms and in constructing DNA based logic circuits (Khodakov et al. 2015; Ravan et al. 2020). The specificity of TMSDRs lie in nucleic acid sequence dependency. TMSDRs are often more sensitive to base-pair mismatches than some of the other classical hybridization reactions (Duose et al. 2012), making it appropriate for detection of specific RNA and DNA sequences.

Hammerhead ribozymes (HHRs) are non-proteinaceous catalytic RNA motifs that can catalyze transesterification reactions (Hammann et al. 2012; Scott et al. 2013). HHRs can be found in plant RNA viruses, satellite RNA, viroids and repetitive satellite DNA (Hammann et al. 2012). HHRs require a minimum of ~50 nucleotides to form a three-dimensional structure resembling a hammerhead and that can catalyze strand scission reactions.(Amarzguioui and Prydz 1998). Synthetic HHRs are capable of cleaving RNA strands *in trans* however, known HHRs also demonstrate *cis* activity (Scott et al. 2013). In nature, there exists (at least) three types of HHRs depending on the positions of loops (Hammann et al. 2012). For example, type I HHR has loops in both Stems II and III, but not in Stem I, as shown in Fig 1. HHRs can be designed to be induced by single-stranded RNA (ssRNA) molecules, single-stranded DNA (ssDNA) molecules or, in case of aptazymes, by small molecules such as tetracycline (Penchovsky and Breaker 2005; Wittmann and Suess 2011). Using these HHRs, different types of logic gates can (and have been) be designed, activated by one or more inputs including ssDNA and ssRNA (Penchovsky and Breaker 2005; Wittmann and Suess 2011).

**FIGURE 1.**
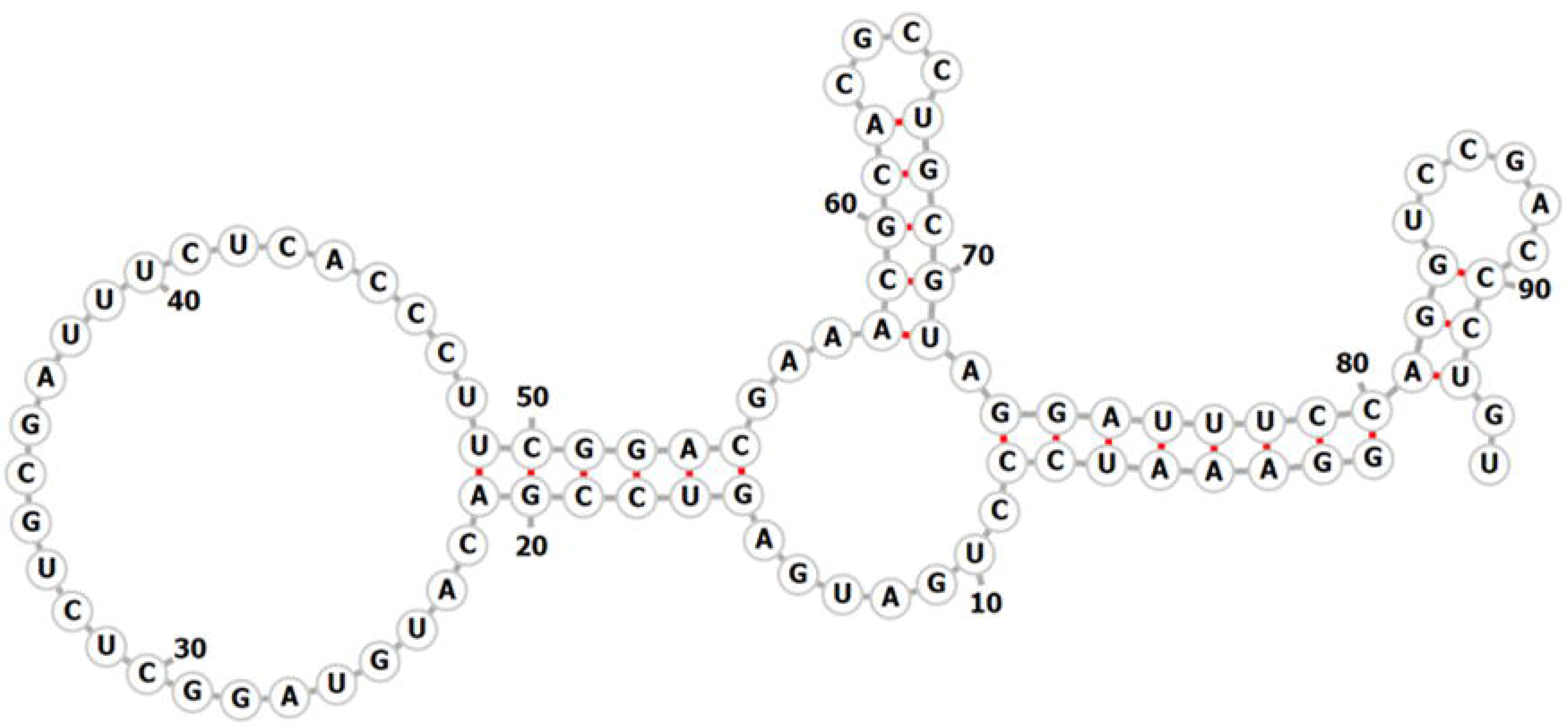
Diagram of the type I hammerhead ribozyme. The black scissors mark the cleavage site.

As illustrated in Fig 2A, a YES logic gate can be described as a single input gate, which is activated upon introduction of input DNA in the medium. In this study, HHRs were designed to respond to specific nucleic acid sequences (inputs). Upon binding with such inputs, these HHRs refold into their active form. This active form catalyzes a strand-cleavage reaction, leading to the production of a small ssRNA fragment called ‘output’. Our designed ribozyme indeed behaved as YES logic gate, as shown by the polyacrylamide gel (Fig 2C).

**FIGURE 2.**
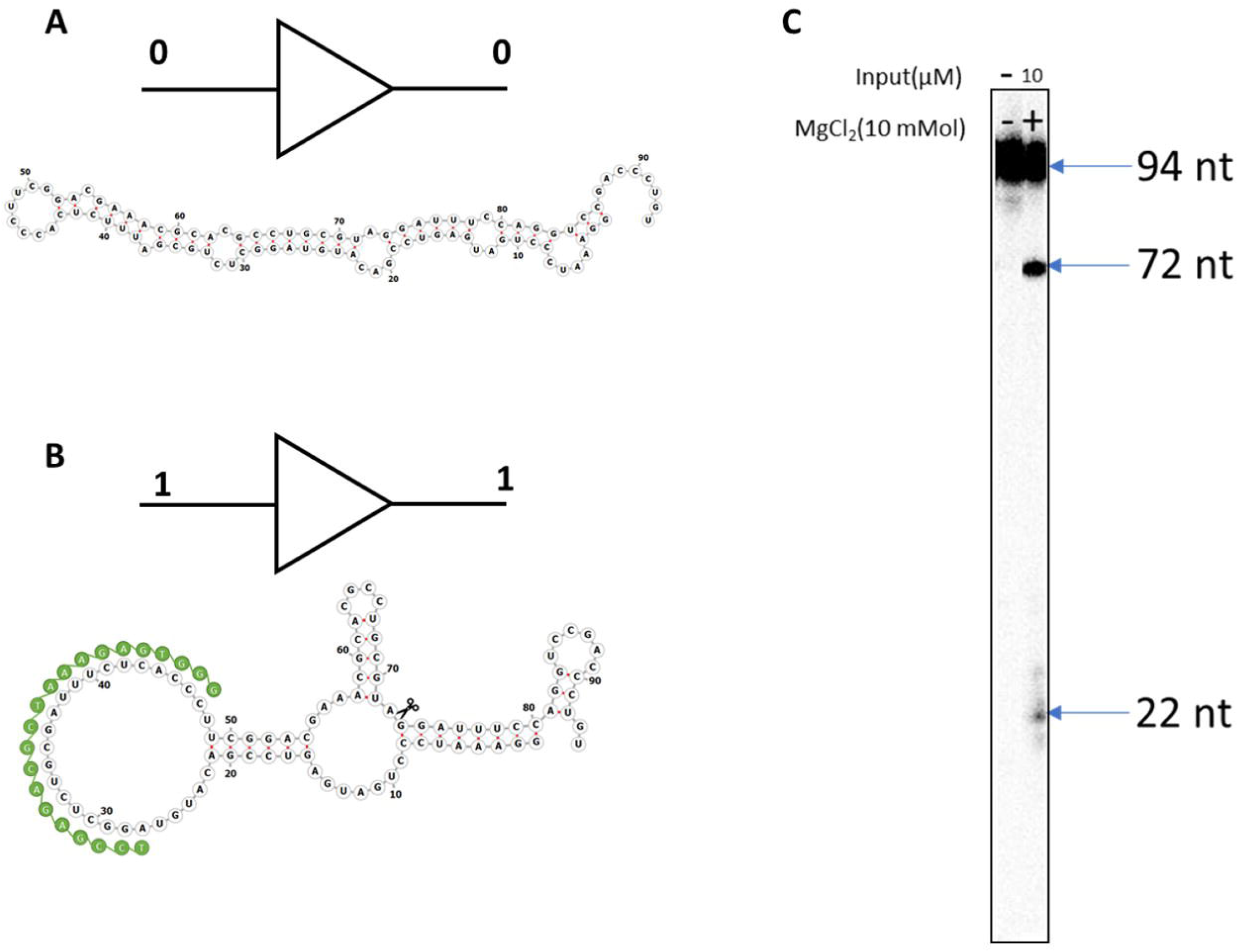
HHR designed to act as YES logic gate. **(A)** Schematic diagram of a YES logic gate without input, and a corresponding inactive HHR without input nucleic acid strand. **(B)** Schematic diagram of a YES logic gate with input, and a corresponding active HHR in the presence of the input DNA oligonucleotide (green strand); the cleavage site is marked by scissors. **(C)** Gel image of YES gate: left lane shows ribozyme without input and MgCl_2_; right lane shows ribozyme with input and MgCl_2_; right lane shows cleavage of ribozyme and expected band size. All ribozyme secondary structures were generated using Forna-RNA webserver (Kerpedjiev et al. 2015).

Evaluating the kinetics of ribozyme cleavage reactions is essential to understanding their mechanisms of operation, for characterizing mutants and for estimating certain parameters-critical to proper functioning of RNA-based circuits, such as cleavage rate. Indeed, ribozymes that appear to have similar activity, based on end-point cleavage, can have orders of magnitude of differences between them, in terms of cleavage reaction rates. Generally, ribozyme kinetics are monitored using a radioisotope of ^32^P that can be incorporated during transcription, or post transcriptionally using a kinase enzyme (Penchovsky and Breaker 2005; Wittmann and Suess 2011; Hammann et al. 2012). In addition to certain disadvantages, such as limited half-life and radiation hazard, the use of radioisotopes for this procedure limits it to laboratories with appropriate facilities. These facilities that are becoming less common since fluorescence is increasingly replacing radioactivity as a preferred method of labeling. Other groups (e.g., (Li et al. 2005)) demonstrated HHR kinetics using fluorescence, where RNA molecules were labeled with cyanine-AMP during transcription. However, this process entails RNA modifications, causing fluctuations in the annealing temperature, which affect RNA folding and hence, ribozyme activity (Li et al. 2005; Scott et al. 2013; Moreira et al. 2015).

In this study, we describe a novel approach to detecting HHR cleavage using the toehold mediated strand displacement reaction (TMSDR). A previously characterized YES logic gate is used to investigate the functionality of the novel TMSDR method (Fig 2). Upon binding to the input strand, the inactive ribozyme (Fig 3A) folds into its active conformation (Fig 3B), leading to HHR self-cleavage and to the release of a small 22nt ssRNA ‘output strand’ (Fig 3C, bottom). When this output strand binds to the fluorescent strand of a specifically designed probe (Fig 3, D and E), the quencher strand is displaced, leaving the fluorophore free to fluoresce (Fig 3F, bottom). The concentration of the cleaved-off RNA fragments dissociated from the ribozyme is determined using a standard curve.

**FIGURE 3.**
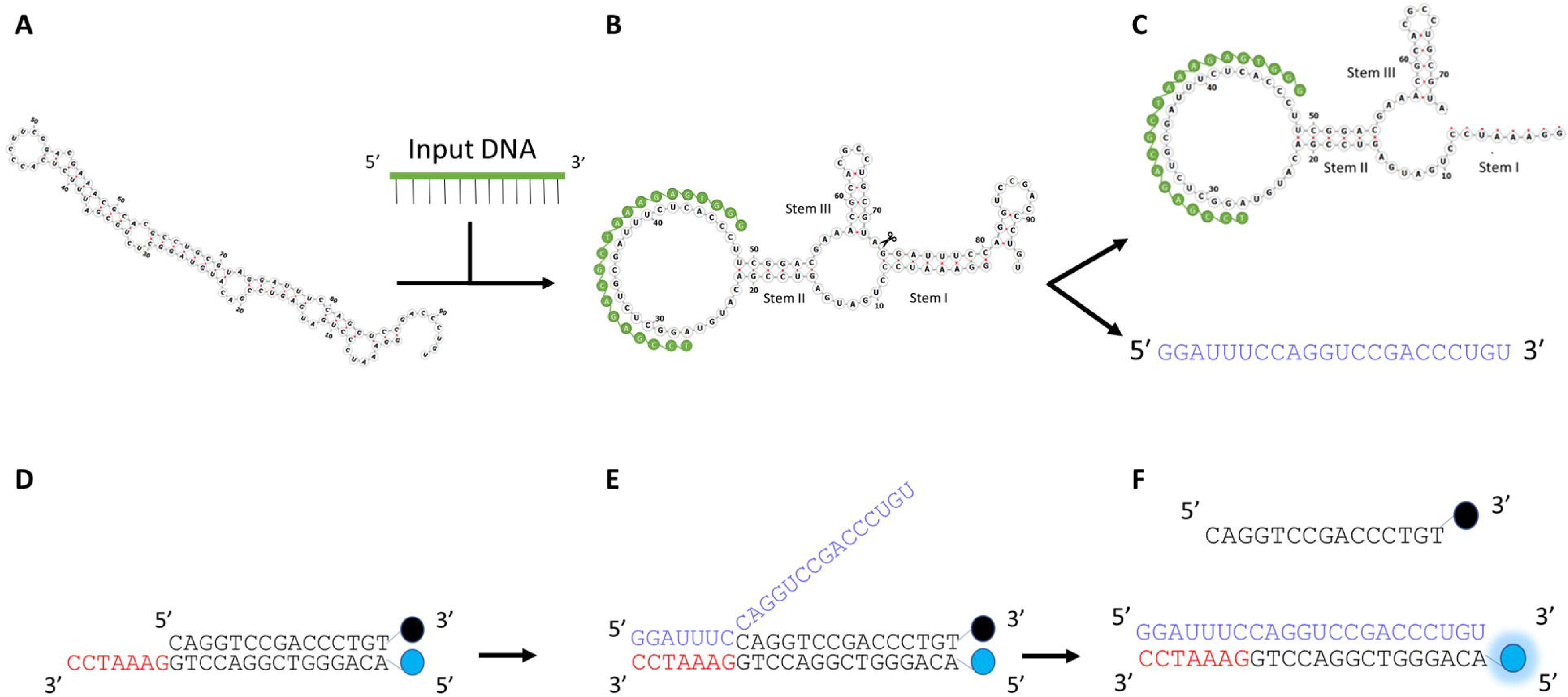
Schematic representation of the toehold mediated strand displacement reaction used in measuring the kinetics of hammerhead ribozyme cleavage reactions. **(A)** Misfolded HHR in absence of input. **(B)** The introduction of input DNA oligo (green strand) induces the formation of an active HHR core, resulting in cleavage activity **(C)** Cleavage products: cleaved HHR bound to input and released output (blue strand) **(D and E)** Released output interacting with toehold present on pre-annealed probe; this interaction results in the displacement of the Q-strand (orange strand) **(F)** Displacement of the Q-strand results in the separation of the quencher from the fluorophore, culminating in detectable fluorescence. All ribozyme figures were generated using Forna-RNA web server (Kerpedjiev et al. 2015).

## MATERIALS AND METHODS

### Ribozyme sequence selection and generation

The algorithm used in this work is an extension of the one in (Penchovsky and Breaker 2005). The main difference is that rather than using random search, the employed algorithm implements an *evolutionary algorithm* (EA) to search for hammerhead ribozymes that function as Yes logic gates. An EA is a search and optimization method that simulates evolution within a computational environment. It operates on a population of candidate solutions (individuals) to a problem. Generally, an EA terminates when one or more satisfactory solutions to the problem are found, stagnation occurs or, an upper limit to computational resources is reached.

### PCR assembly of DNA template

Predesigned overlapping oligodeoxynucleotides (Fig S1) were assembled by PCR (BioRAD T100) using Primers F1, R1, F2 and R2 (Fig S1 and Table S1). The PCR reaction was carried out in a fixed volume of 100 μl, containing primers F1 (2 μM), R1 (0.2 μM), F2 (0.2 μM), R2 (2 μM), Taq polymerase (hotStar Taq Plus from QIAGEN) with its reaction buffer at 1x, Q-solution (1x from QIAGEN), 0.2 mM of dNTPs (DGel electrosystem) and miliQ water. The reaction mixture was subjected to 15 min denaturation at 95°C and 15 cycles consisting of: 30 s denaturation at 95°C, 30 s annealing at 50°C and 30 s extension at 72°C. PCR was validated by visualizing 5 μl of reaction mixture on 2% agarose gel containing gel red (Trans). The remaining PCR product was ethanol precipitated.

### In vitro transcription and RNA purification

In vitro RNA synthesis was performed as previously described (Perreault et al. 2011), with slight modifications. When larger quantities were required, the reaction was carried out in a fixed volume of 1 ml. The reaction mixture contained 80 mM HEPES (pH 7.5), 24 mM MgCl_2_, 40 mM dithiothreitol, 2 mM spermidine, 6μg/ml T7 polymerase, 150 μl of PCR product (For 1ml transcription, 10 PCR reactions (100 μl each) were pooled together, precipitated and resuspended in 150 μl miliQ water), 2 mM rNTPs, 1x pyrophosphatase (Roche diagnostics) and 200U (40U/μl) RiboLock (Thermo Fisher Scientific). The reaction mixture was incubated at 37°C for 150 minutes, treated with 10U of DNase (New England Biolabs), incubated at 37°C for 30 minutes. The RNA was extracted with phenol-chloroform, and the aqueous phase was ethanol precipitated. The RNA was purified in 10 % denaturing (8 M urea) polyacrylamide gel. The gel was revealed by UV-shadowing. The band of interest (Highest band on gel, as there was some level of cleavage during transcription) was excised and eluted in 0.3 M NaCl overnight at 4°C. The eluent was ethanol precipitated and resuspended in nuclease free water.

### Radiolabeling of ribozyme using [α-^32^P] UTP during transcription

Radiolabeling of RNA was conducted as previously described with minor modifications. Here, the reaction mixture consisted of 1X transcription buffer (see above), 15 μl of PCR product (100 μl PCR reaction ethanol precipitated and resuspended in 20 μl miliQ water), 2 mM of GTP,CTP,ATP, 0.125 mM UTP, 1x pyrophosphatase (Roche diagnostics) and 40 U RiboLock (Thermo Fisher Scientific) and 1 μl of [α-^32^P] UTP (Perkin Elmer) per 50 μl reaction. The reaction mixture was ethanol precipitated and analyzed in 10% denaturing polyacrylamide gel; the product was revealed by phosphorimaging (Typhoon 9500 FLA; GE Healthcare Life Sciences). The band of interest was resected and eluted in 0.3 M NaCl overnight at 4°C. The eluent was ethanol precipitated and resuspended in nuclease free water.

### Preparation of fluorescent probe

Pre-designed oligodeoxynucleotides were conjugated at the 5’-end with Cy5 and at the 3’-ends with Black hole quencher (Alpha DNA, Montreal, Canada). The strand with the Cy-5 at the 5’-end was named the ‘F-strand’ (5’-ACAGGGTCGGACCTGGAAATCC-3’), while the strand with the Black hole quencher-3 (BHQ-3) at 3’-end was called the ‘Q-strand’ (5’-CAGGTCCGACCCTGT-3’) (Fig 3D). The probe was prepared in a cleavage buffer (100 mM NaCl, 50m M tris-HCl pH 7.5, 25 mM KCl) with 0.5 μM F-strand and 0.6 μM Q-strand per 10μl reaction. The reaction was incubated in thermocycler (BioRad T100) for 3 min denaturation at 95° C, 15 min annealing at 50° C and 15 min annealing at 37° C.

### Calibration of probe and standard curve generation

The prepared probe was calibrated using ssRNA oligonucleotide (IDT) mimicking the ribozyme output (5’-GGAUUUCCAGGUCCGACCCUGU-3’). We called this strand the ‘R-strand’ (Displacer RNA-strand). Different concentrations of R-strand, ranging from 0.05 μM to 2 μM were mixed with 0.5 μM preannealed probe. The reaction mixture was incubated at 37° C and analyzed using a fluorescent plate reader (Tecan M1000 pro) at 647 nm excitation and 665 nm emission wavelengths. The probe was also calibrated using DNA displacer strand called ‘D-strand’. Comparison of R-strand and D-strand standard curve is illustrated in Fig S4.

### Analysis of Hammerhead ribozyme kinetics on polyacrylamide gel

Ribozyme kinetics were assayed using a prelabeled [α-^32^P] UTP ribozyme. The reaction was performed in a fixed volume of 10μl, containing 100 mM NaCl, 50 mM Tris-HCl pH-7.5, 25 mM KCl, 10 mM MgCl_2_, 10 μM input oligodeoxynucleotide and 1 μl of the labelled ribozyme. The reaction was started by adding MgCl_2_. The reaction was incubated at 37 °C. Sequentially, the aliquots of reactions reaction were stopped at 30 minutes intervals using denaturation buffer (80 % formamide, 0.5 mM EDTA, 0.02% bromophenol blue and 0.02% xylene cyanole). The samples were analyzed on 10% denaturing polyacrylamide gel, the gel was developed by phosphorimaging and the band intensity was determined using ImageQuant software (GE Healthcare Life Sciences).

### Analysis of hammerhead ribozyme kinetics with strand displacement

A preannealed probe was used to evaluate HHR cleavage kinetics. Here, 0.5 μM of a preannealed probe was mixed with 10 mM MgCl_2_, 10 μM input oligodeoxynucleotide and 1 μM ribozyme per 10 μl reaction. The reaction mixture was incubated at 37°C. The fluorescence emitted was measured using a fluorescent plate reader (Tecan M1000 Pro). Readings were taken every 30 minutes.

## RESULTS

### Probe design and workflow of probe mediated hammerhead kinetics

The ribozyme is designed to be *inactive* by default, then turn *active* when bound to an input strand. The secondary structure of the inactive form of the ribozyme is shown in Fig 3A. Once the input strand is bound to the oligonucleotide biding site (OBS) of the ribozyme, the ribozyme refolds into an active conformation (Fig 3B) causing self-cleavage. Ribozyme self-cleavage releases the output strand (Fig 3C), which binds progressively to the toehold of the probe (Fig 3, E and F). A few assays (data not shown) revealed that a toehold of 7nt is sufficient to displace the quenching Q-strand. The probe is designed so strand displacement proceeds from the 5’-end to the 3’-end of the Q-strand, as this has been established to be most efficient route to displacement (Šulc et al. 2014; Simmel et al. 2019). At the conclusion of the strand-displacement reaction, the quencher fully separates from the fluorophore, allowing the Cy5 to fluoresce.

### Kinetics of ribozyme cleavage with radiolabeled RNA

From previous studies, it has been shown that the radiolabeling of RNA molecules constitutes one of the best approaches to investigating RNA structure and function, *in vitro* (Celander and Cech 1991; Sclavi et al. 1998; Li et al. 2005). However, the process of radiolabeling RNAs is a laborious one, requiring the utilization of radioisotopes, which many labs cannot handle, and specialized analysis equipment (Porecha and Herschlag 2013).

To investigate the cleavage of HHRs in the presence of inputs, HHRs are labelled with [α-^32^P] UTP during transcription, incubated in a cleavage buffer, and the reaction is sampled (and stopped) at different time-points, to provide inputs to a polyacrylamide gel display. To provide further evidence in support of the claim, the original ribozyme (Fig 1) was modified at the 5’-end to increase base pairing between the expected output strand and its complementary strand in stem I. Two of those ribozymes were designed (Fig 4, A and B), and then assayed alongside the original ribozyme, using the radiolabeling approach. The completed gel exhibits comprehensive data on the temporal progress of the cleavage reactions. In Fig S3, the two major bands of each gel image correspond to the size of the full-length ribozyme and its cleaved product. For the original ribozyme the two bands indicate 94nt and 72nt, respectively; for Ribozyme+2bp they indicate 96nt and 74nt, respectively; for Ribozyme+14bp they indicate 108nt and 86nt, respectively. An increase in band intensity with time was measured for the cleavage products of all three ribozymes (also shown in Fig S3). The intensity of these bands were used as a measure of percentage of cleaved ribozyme: this is equal to the ratio of cleaved product (Cleaved ribozyme + Output fragment) over cleaved product plus full-length ribozyme (Uncleaved ribozyme + Cleaved ribozyme + Output fragment).

**FIGURE 4.**
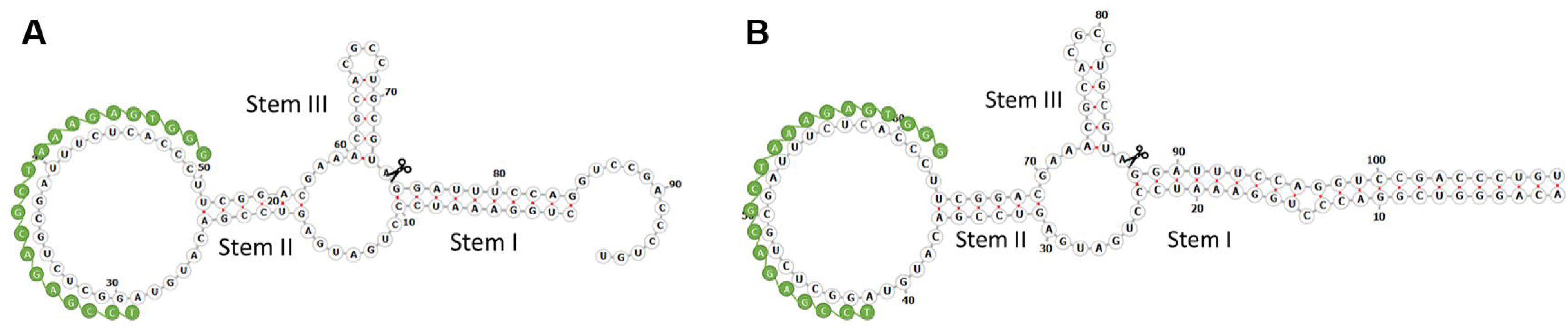
Different active HHR secondary structures, with different amounts of base-pairing in stem I between the still-attached output strand and its complement. The green strand represents the input DNA oligonucleotide for all ribozymes (5’-GGGTGAGAAATCGCAGAGCCTA-3’). **(A)** A YES logic gate with 10nt of base-pairing in stem I (Ribozyme + 2 bp); **(B)** a YES logic gate with 22nt of base-pairing in stem I (Ribozyme+14bp). Both ribozyme images were generated using Forna-RNA webserver (Kerpedjiev et al. 2015).

As expected, there was a gradual increase of cleavage percentage with time, for all three ribozymes (Fig 5). These results indicate that the input sequence does indeed induce HHR cleavage, under typical cleavage conditions (i.e., in presence of Mg^2+^). This shows that the designed HHRs have functioned as YES logic gates, in response to their intended input (Fig 2C and Fig S2). As an unintended result, we also found that the cleavage efficiencies of the different ribozymes were markedly different (Fig 5). We hypothesize that this was due to different interactions between the upaired segment of stem I and the rest of the ribozyme; testing this hypothesis and charaterizing the exact nature of such possible interactions fall outside the scope of this study.

**FIGURE 5.**
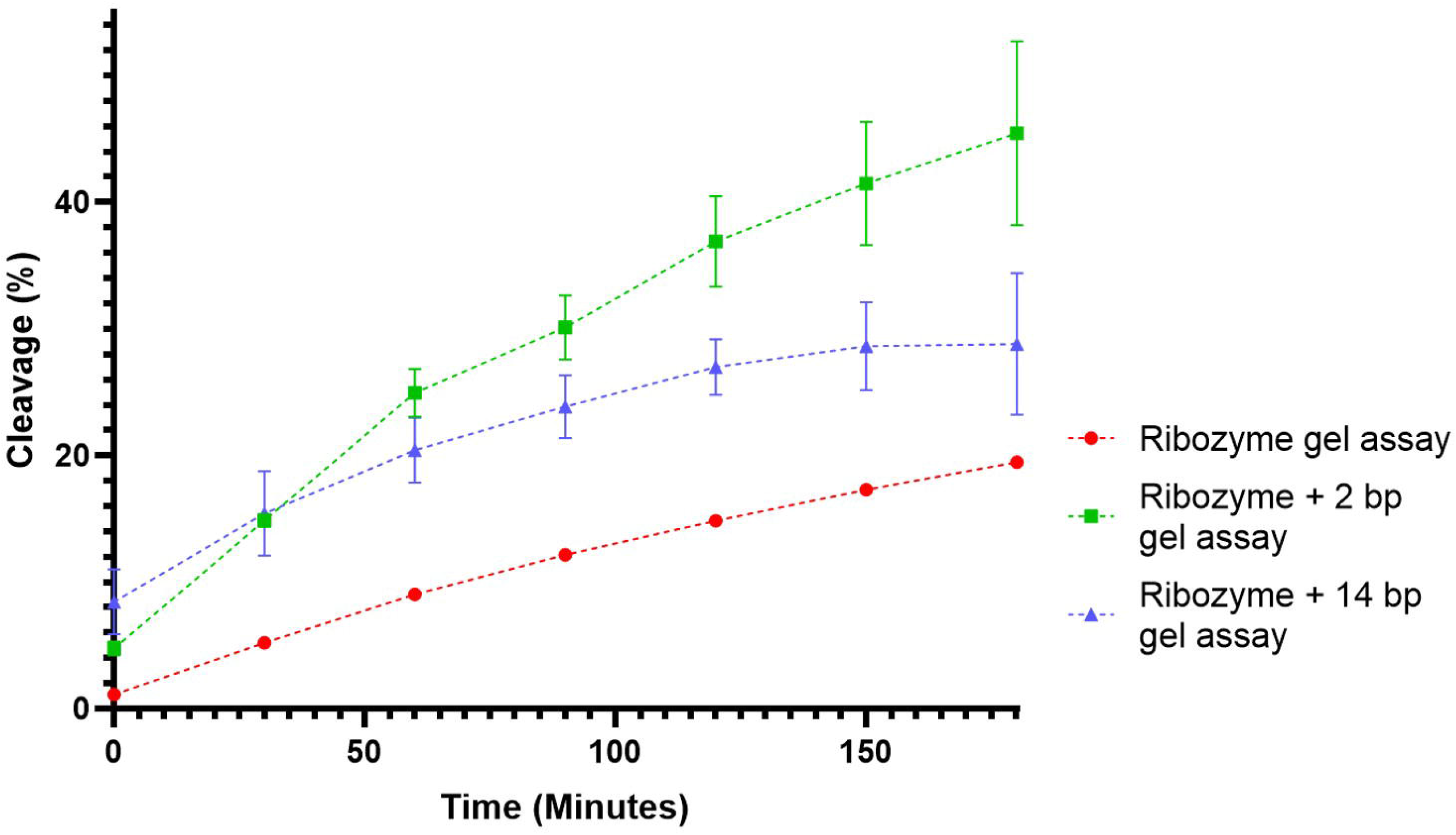
HHR kinetics using [α-^32^P] UTP labelling. Standard cleavage buffer conditions were used, and all the reactions were carried out at 37°C. All cleavage reactions were performed in triplicates.

### Evaluation of HHR kinetics via a strand displacement reaction utilizing a fluorescent probe

The assessment of HHR kinetics using radiolabeling suffers from certain limitations. For example, determining the concentrations of cleaved ribozymes and cleaved detached RNA outputs using denaturing gels is problematic. This is because a gel shows all output strands of equal lengthin the same band, whether these strands have actually detached from the rest of the ribozyme or not, post cleavage. In addition, this method is time consuming and involves the use of radioisotopes, which are carcinogenic (Furth and Tullis 1956). To overcome these limitations, we sought to evaluate HHR kinetics using predesigned fluorescent probes.

Interestingly, we noticed (Fig 6A) an increase in fluorescence intensity with time in the assay group (HHR with input and Mg^2+^). However, little or no change in fluorescence intensity was noticed in either the background group or the HHR alone group (Fig 6A). Interestingly, for both Ribozyme+2bp and Ribozyme+14bp assay groups, the observed fluorescence was near background levels. Taken together, these results provide evidence that the cleaved output from the original ribozyme binds to the toehold, displaces the Q-strand, leading to the observed fluorescence.

**FIGURE 6.**
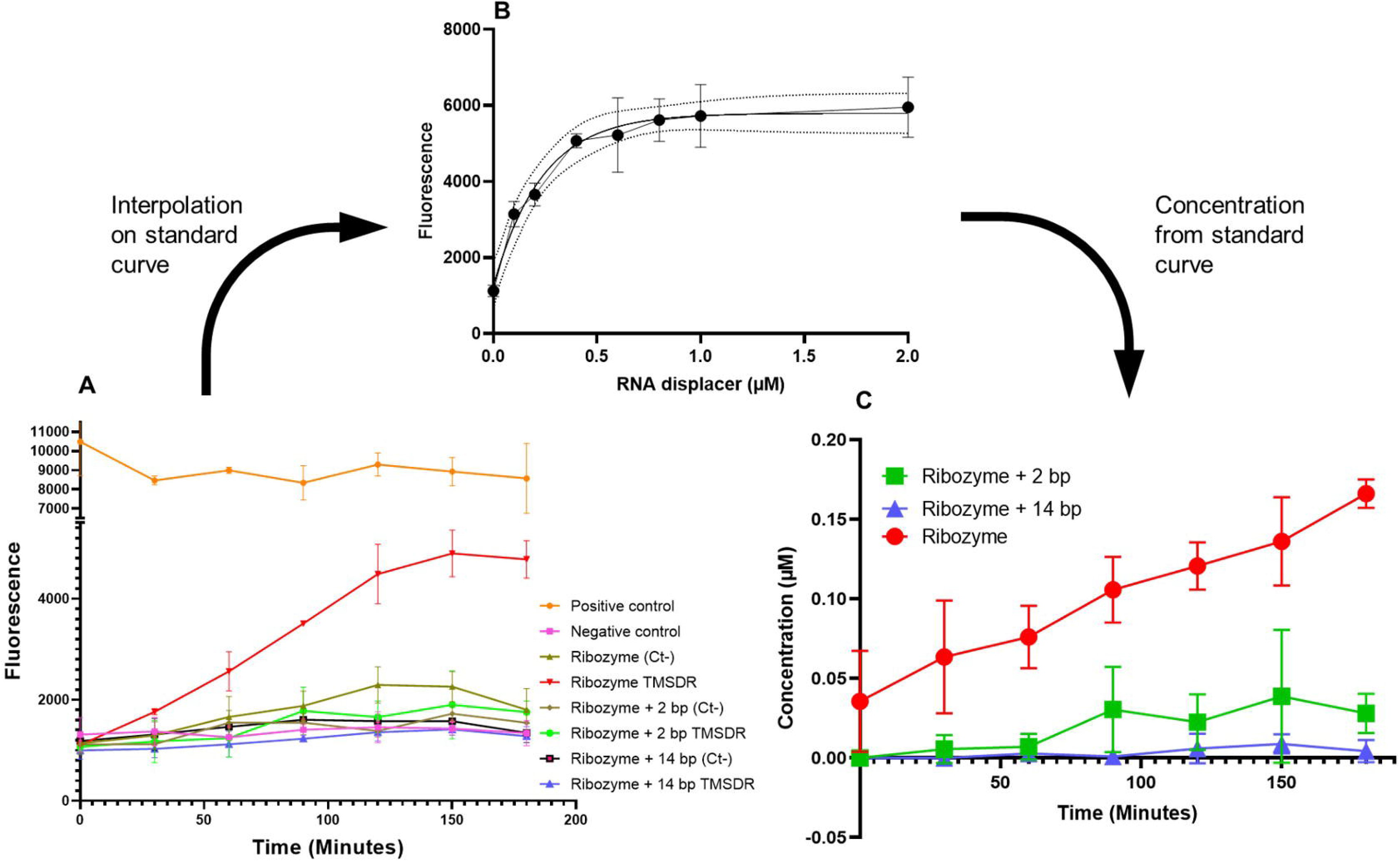
Analysis of the YES gate using probe (with Cy-5 as fluorophore and Black hole quencher as quencher). **(A)** 0.5μM of probe and 1μM of ribozyme were used in the assay. 10μM of the R-strand with the probe was used as positive control, and a quenched probe was used as negative control. Ribozyme without Mg^2+^ and without input DNA was used as another negative control (Ct). The assay group includes 10μM input DNA, 10 mM Mg^2+^. Readings were taken every 30 minutes over a period of 180 minutes. The same protocol was followed for Ribozyme+2bp and for Ribozyme+14bp. **(B)** The standard curve for the 0.5μM probe using the same reagents as for the assay. Different concentrations of the RNA displacer strand were used (0.05, 0.1, 0.2, 0.4, 0.6, 0.8, 1 and 2 μM). **(C)** Interpolated fluorescence values from the three ribozyme assays mapped using a standard curve to concentrations of released output strand, for the same three ribozymes.

Furthermore, to determine the concentration of HHR output, we generated a standard curve using an R-strand equivalent to the output strand (Fig 6B). Different concentrations of R-strands were mixed in with the probe and assayed using a fluorescent plate reader. We observe a stoichiometric relationship between R-strand concentration and fluorescence (Fig 6B). These results demonstrate an increase in TMSDR fluorescence as a function of increased R-strand concentration. Thus, the generated standard curve can be utilized to interpolate the fluorescence values obtained from the TMSDR assay and hence, determine the concentration of detached output strand, generated by ribozyme self-cleavage.

Interpolated values were plotted for all three ribozymes. The original ribozyme shows the highest activity level (as determined by TMSDR) relative to ribozyme+2bp and Ribozyme+14bp (Fig 6C). The original ribozyme has only 8 base pairs in stem I, joining the output strand to its complementary strand (Fig 1). Two mutant ribozymes were generated: a ribozyme with 10 base pairs in stem I (Ribozyme+2bp) and another ribozyme with 22 base pairs in stem I (Ribozyme+14bp) (Fig 3A and B). The TMSDR results (Fig 6A, further highlighted in Fig S5) show a decrease in fluorescence as a function of increased base-pairing with the output strand.

### Comparison of measurement of HHR cleavage reactions using conventional gels versus TMSDR

To better evaluate the output concentration derived from the conventional approach (gel) and the new probe approach (TMSDR), we compared cleavage activity measured by gel band intensity with cleavage activity as reflected by probe fluorescence (Fig 7A). These results indicated that the two approaches measure the progress of cleavage reactions in different ways and provide complementary information: breakage of the phosphodiester linkage at the cleavage site measured with the denaturing gel vs. amount of dissociated products measured by TMSDR.

**FIGURE 7.**
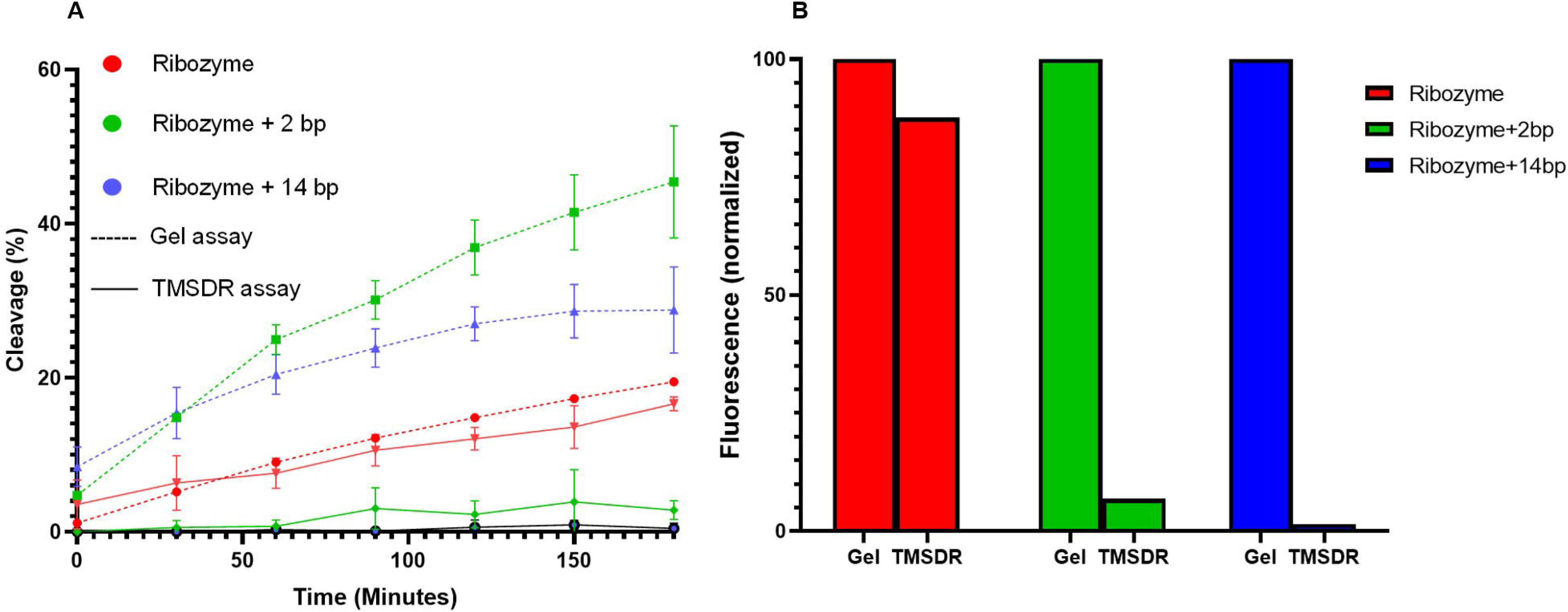
TMSDR vs. Gel cleavage analysis **(A)** Comparison of cleavage obtained from [α-^32^P] UTP labelled ribozymes (dotted lines) and TMSDR (solid lines). **(B)** The area under the curve, representing total emitted fluorescence (normalized), was calculated from the graph for the original ribozyme, ribozyme+2bp and ribozyme+14bp.

As demonstrated in Fig 7B, the cleavage from the gel is comparable with cleavage derived from the TMSDR assay in case of original ribozyme (normalized). However, as the base pairing with the output strand increases, even by as little as 2nt, the amount of released output decreases considerably, as illustrated by the green bars representing ribozyme+2bp. In case of ribozyme+14bp, virtually no cleavage activity was observed from TMSDR as compared to its gel counterpart. It is important that the experimenters understand that and hence, utilizes the method most appropriate to the particular needs of their own projects.

## DISCUSSION

In the past decade, several methods have been developed and used to analyze and evaluate the structure, function and activity of ribozymes *in vitro*. These methods include RNA radiolabeling, post-transcriptional fluorescence labelling, phosphoramidite chemistry for fluorescent labelling and engineered fluorescent aptamers (e.g., Spinach and Mango) (Li et al. 2005; Porecha and Herschlag 2013; Auslander et al. 2016; Mitra and Ha 2019; Debiais et al. 2020). These techniques make use of transcriptional incorporation of [α-^32^P] UTP, 5 ‘-incorporation of ^32^P from [γ-^32^P] ATP or fluorophore, chemical synthesis of RNA and fluorescence activity of aptamers (Singh et al. 1999; Li et al. 2005; Porecha and Herschlag 2013; Auslander et al. 2016). However, besides radiolabeling, which has its own disadvantages, these methods are associated with direct RNA modification, which in turn can impact the structure, function and thermodynamic stability of the measured ribozyme (Li et al. 2005; Moreira et al. 2005).

In this study, we present a novel approach for hammerhead ribozyme (HHR) cleavage kinetics that utilizes the toehold the mediated strand displacement reaction (TMSDR). The proposed method separates the detection system from the ribozyme, eliminating the need for ribozyme labeling and modification. This fosters unhindered determination of ribozyme kinetics.

We conceived an oligonucleotide activated HHR, which functions as a YES logic gate. When the HHR binds to the input oligonucleotide, the HHR cleaves itself, generating an ssRNA fragment that can detach from the HHR. Detached output ssRNA interacts with the toehold present on (the 3’-end of the F-strand) of a dsDNA probe. This binding initiates a strand displacement reaction favoring the expulsion of the (Q-strand) quencher in a 5’ to 3’ direction. This process culminates in the dissociation of the quencher from the fluorophore, resulting in detectable fluorescence.

The vast majority of previous studies determine HHR cleavage using radiolabeling or product separation (Penchovsky and Breaker 2005; Perreault et al. 2011; Hammann et al. 2012; Porecha and Herschlag 2013). Product separation on denaturing gels is associated with forced detachment of an output strand from an HHR. Consequently, this approach fails to distinguish between released output and cleaved but bound output. However, in TMSDR, the occurrence of fluorescence is a direct measure of released output. A gradual increase in cleavage activity over time was noted in both (gel and TMSDR) methods for the ribozyme. However, a decrease in cleavage activity from TMSDR was observed in case of ribozyme+2bp and ribozyme+14bp, compared to their gel counterparts. This decrease in activity is correlated with an increase in the strength of binding between the cleaved output strand and its complement (on stem I) of the ribozyme. This decreased activity is indicative that TMSDR-based fluorescence is a measure of the concentration of the *detached* output strand, rather than the full extent of ribozymatic self-cleavage. Thus, TMSDR allows for real-time cleavage monitoring and realistic evaluation of the amount of product (RNA output) leaving the ribozyme, rather than mere cleavage.

Therefore, our approach allows for measurement of released output strands, which incidentally is more important than cleavage only for many synthetic biology applications. Furthermore, when complementing traditional radiolabeling methods, a TMSDR-based technique can help provide a more complete picture of cleavage activity and rate of dissociation of the cleaved-products. This information that can be crucial to determining and characterizing the limiting step for the development of ribozyme-based RNA circuits. TMSDR lends itself to automation more readily than radiolabeling-a trait particularly useful for eventual design of more complex RNA logic gates and circuits.

## ACKNOWLEDGEMENTS

The authors wish to thank Ms. Sabrine Najeh for technical assistance and helpful discussion.

## FUNDING

Mr. J. Kapadia and Mr. N. Kamel received fellowships from Natural Sciences and Engineering Council of Canada (NSERC) SynBioApps program [CREATE-511601-2018]. J.P. is a junior 2 FRQS research scholar. Other funding, including for open access charge: NSERC [RGPIN-2019-06403].

